# Elaboration of a MALDI-TOF Mass Spectrometry-based Assay of Parkin Activity and High-Throughput screening platform for Parkin Activators

**DOI:** 10.1101/2022.03.04.482851

**Authors:** Ryan Traynor, Jennifer Moran, Michael Stevens, Axel Knebel, Bahareh Behrouz, Kalpana Merchant, C. James Hastie, Paul Davies, Miratul M. K. Muqit, Virginia De Cesare

**Affiliations:** MRC Protein Phosphorylation and Ubiquitylation Unit, School of Life Sciences, University of Dundee, Dow St, Dundee DD1 5EH, Scotland, UK; Vincere Biosciences Inc, 245 Main St. Fl 2, Cambridge, MA 02142, USA; MRC Protein Phosphorylation and Ubiquitylation Unit Reagents and Services, School of Life Sciences, University of Dundee, Dow St, Dundee DD1 5EH, Scotland, UK

## Abstract

Parkinson’s disease (PD) is a progressive neurological disorder that manifests clinically as alterations in movement (bradykinesia, postural instability, loss of balance, and resting tremors) as well as multiple non-motor symptoms including but not limited to cognitive and autonomic abnormalities. Mitochondrial dysfunction has been linked to sporadic PD and loss-of-function mutations in genes encoding the ubiquitin E3 ligase Parkin and protein kinase, PTEN-induced kinase 1 (PINK1), that regulate mitophagy, are causal for familial and juvenile PD^1-3^. Among several therapeutic approaches being explored to treat or improve PD patient’s prognosis, the use of small molecules able to reinstate or boost Parkin activity represents a potential pharmacological treatment strategy^4^. A major barrier is the lack of high throughput platforms based on robust and accurate quantification of Parkin activity *in vitro*. Here we present two different and complementary Matrix Assisted Laser Desorption/Ionization-Time of Flight mass spectrometry (MALDI-TOF MS) based approaches for the quantification of Parkin E3 ligase activity *in vitro*. These methods recapitulate distinct aspects of ubiquitin conjugation: Parkin auto-ubiquitylation and Parkin-catalysed discharge on lysine residues. Both approaches are scalable for high-throughput primary screening to facilitate the identification of Parkin modulators.

## Introduction

The coordinate action of the RING-IBR-RING (RBR) E3 ubiquitin ligase Parkin and PTEN-induced kinase 1 (PINK1) is fundamental for the clearance of dysfunctional mitochondria by mitophagy in nearly every cell type including neurons ^5-7^. Under healthy cellular conditions, Parkin is present in the cytosol in an auto-inhibited conformation^8-10^. Upon mitochondrial depolarisation, that can be induced by mitochondrial uncouplers, PINK1 is activated and recruits and activates Parkin at sites of mitochondrial damage via directly phosphorylating Parkin at Serine 65 within its Ubiquitin-like domain (p-Parkin), and indirectly by phosphorylating ubiquitin (p-Ub) also on Serine 65^11-13^. Under *in vitro* assay conditions of Parkin activation, Parkin and p-Parkin undergo auto-ubiquitylation on accessible lysine residues; can catalyse ubiquitin transfer to substrates; or can stimulate discharge of ubiquitin from charged E2 enzyme, UBE2L3, onto primary amines present in the reaction buffer (discharge assay). Both types of ubiquitylation have previously been used as read-outs for Parkin and p-Parkin activity determination^13^. Low-throughput, SDS page based-technique have been extensively applied for visualizing Parkin autoubiquitylation patterns^13,14^. Ubiquitin-fluorescent probes have also been developed to exploit the reactivity of Parkin toward primary amines^15^. While both these approaches have proved to be easy tools for investigating Parkin activity *in vitro*, they have substantial caveats and limitations. SDS-page based techniques lack scalability to high-throughput format and are often not fully quantitative while fluorescent based-approaches are intrinsically prone to fluorescence related artefacts. Herein we have elaborated two quantitative and complementary Matrix Assisted Laser Desorption/Ionization-Time of Flight mass spectrometry (MALDI-TOF MS) based assays to determine Parkin and p-Parkin activity *in vitro*. Both methods allow for quantitative investigation of Parkin activity *in vitro*; are scalable to high-throughput formats and employ physiological substrates (ubiquitin and p-ubiquitin) thus circumventing artefacts associated with the use of fluorescence-based tools.

## Results

### Development of MALDI-TOF Parkin Activity assays

We previously reported the development of a MALDI-TOF MS based method for the quantification of the activity of ubiquitin E2 conjugating enzymes and E3 ligases belonging the RING, HECT and RBR family^16^. While RING E3 ligases rely on the catalytic activity of a cognate E2 conjugating enzyme, HECT and RBR E3 ligases receive ubiquitin from the E2 conjugating enzymes to ubiquitylate their substrate on lysine residues. Most E3 ligases will undergo autoubiquitylation when tested *in vitro*. The previously published MALDI-TOF E2/E3 assay was based on quantification of the progressive disappearance of ubiquitin as a consequence of its utilisation in the auto-ubiquitylation process^16^. Since the reactivity towards lysine is mediated by the E3 ligase in the RBR enzymatic cascade, we explored whether we could determine Parkin reactivity using a complementary lysine discharge method (also known as nucleophile reactivity assay^17^) of untagged (His-SUMO cleaved) recombinant human Parkin expressed in *E*.*coli* as previously described^18^. UBE2L3 is a HECT-RBR specific E2 conjugating enzyme that lacks intrinsic E3-independent reactivity towards lysine residues^17^. Therefore, the ability to discharge on lysine – and the consequent formation of ubiquitin-lysine adducts (Ub-K) - relies exclusively on the activity of a cognate HECT or RBR E3 ligase. In the auto-ubiquitylation assay, quantification of Parkin activity is achieved by comparing the signal of ubiquitin to that of the heavy-labelled ubiquitin internal standard (^15^N-Ub). Therefore, the autoubiquitylation rate can be represented as a linear reduction of detectable ubiquitin over time (Residual Ubiquitin %) (Fig. 1A). In contrast, in the discharge assay, both substrate (Ub) and product (Ub-K) change over time as the former is converted to the latter. Consequently, the mathematical representation of the discharge assay method will be a non-linear function, as both substrate and product measurements change over time (Fig. 1B). Therefore, in the discharge assay, a dedicated standard curve must be defined in advance to determine the rate of product formation (Ub-K Formation %) (Fig. 1B and Sup Fig 5). The unique regulation of WT Parkin requires the combined use of ubiquitin and phosphorylated ubiquitin (p-Ub). The interaction between p-Ub and Parkin releases Parkin’s autoinhibitory state, therefore p-Ub functions as an allosteric Parkin modulator. Due to the closeness in molecular weight between p-Ub (8646.7 *m/z*) and the ^15^N ubiquitin internal standard (8669.7 *m/z* observed), we employed His_6_-tagged-p-Ub (p-Ub-His, 9812 *m/z*, See Sup. Fig 1) to prevent interference with the ^15^N ubiquitin signal. The His_6_ tag present at the C-terminus of p-Ub-His and the absence of a final glycine dyad (See Sup Fig. 1) do not allow for the incorporation of p-Ub-His into poly-ubiquitin chains, although lysine available in the p-Ub-His may still be employed as ubiquitin substrate. The autoubiquitylation assay exhibited slower kinetics compared to the discharge assay and therefore, to achieve comparable reaction rates, the autoubiquitylation assay was performed at 37 ºC while the discharge assay was performed at room temperature. Due to the difference in the experimental settings, the results obtained from the two assays cannot be directly compared, although they aligned each other very closely.

**Figure 1.**
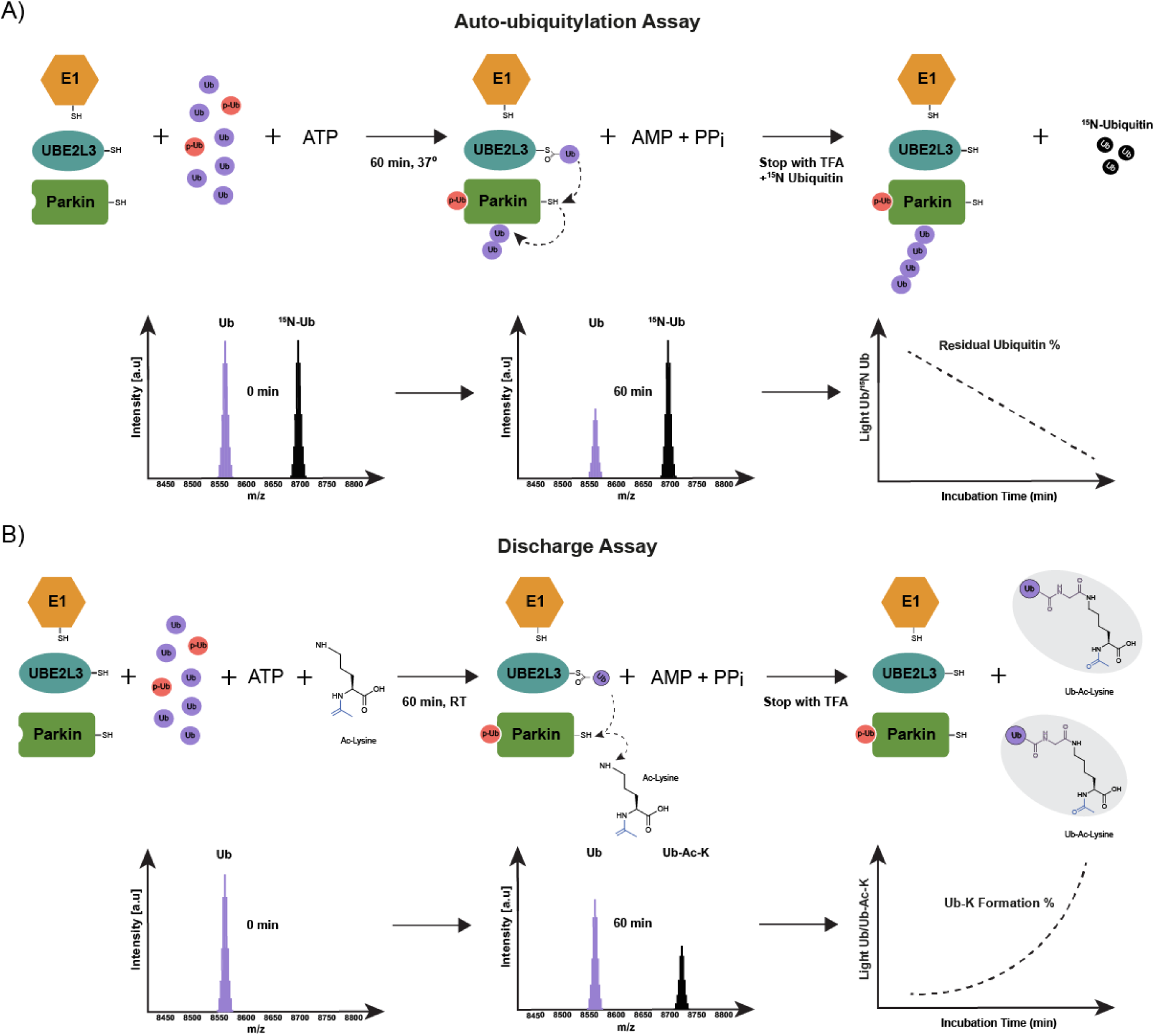
Schematic representation of MALDI-TOF MS Parkin auto-ubiquitylation and discharge assay. A) Parkin auto-ubiquitylation reduces the pool of ubiquitin detected by MALDI-TOF MS over time. Quantification is achieved by use of ^15^N ubiquitin as internal standard consistently present in the reaction (Light Ub/^15^N Ub). B) Parkin-dependent formation of Ub-Ac-Lysine (Ub-K) is detected by MALDI-TOF MS. Quantification is achieved by measuring the ratio between the substrate (Ub) and the product (Ub-K). A linearity curve allows to translate Ub/Ub-K ratio into Ub-K formation %.

### Assessing Parkin activity by MALDI-TOF MS autoubiquitylation and discharge assay

The activity of Parkin is tightly regulated both by direct phosphorylation and by the interaction with phosphorylated ubiquitin^6,13^. We employed the MALDI-TOF autoubiquitylation and discharge assays to accurately quantify the contribution of these regulatory layers on Parkin activity rate. In the autoubiquitylation assay, Parkin activity was quantified by the progressive reduction of the mono-ubiquitin peak (Fig. 1A) while in the discharge assay Parkin activity was assessed by the formation of Ub-Ac-K product (Fig. 2A). Both MALDI-TOF MS methods were employed to quantify the increment in recombinant Parkin and p-Parkin (expressed as previously described^18^, Sup. Fig 3A-B) activity rates upon addition of p-Ub-His. WT Parkin and p-Parkin were tested at a final concentration of 500 nM. Reactions were started by the addition of ubiquitin supplemented with three different concentrations of p-Ubi: 100 nM, 500 nM and 2500 nM. In the autoubiquitylation assay, data were firstly normalized over the ^15^N Ubiquitin internal standard signal (Light Ub/^15^N Ub) and a control reaction without Parkin present (E1+E2 control) was used to establish the rate of Parkin-dependent ubiquitin consumption (Sup. Fig1A and B). We found that an amount of p-Ub-His stoichiometrically equivalent to WT Parkin (500 nM) is sufficient to partially activate WT Parkin (Fig. 2A), while 5 times excess of p-Ub-His induced WT Parkin activity levels comparable to those of p-Parkin in absence of p-Ub-His (Fig. 2A). Stoichiometric amounts of p-Ub-His double the autoubiquitylation rate of p-Parkin after 10 minutes (Residual Ubiquitin 66% in absence of p-Ub compared to 31.5% in presence of 500 nM p-Ub). In the discharge assay, WT Parkin is efficiently activated by stochiometric amounts of p-Ub-His. A similar effect was observed for p-Parkin, whose activity is greatly enhanced already in presence of sub-stochiometric amounts of p-Ub-His (Ub-K Formation 22 % in absence of p-Ub compared to 78% in presence of 100 nM p-Ub) (Figure 2B). Phos-tag SDS-gel analysis indicated that about 80% of Parkin was phosphorylated (Sup Fig. 3), therefore, when testing p-Parkin in presence of p-Ub-His, it is not possible to discriminate whether the increase in activity is due to the activation of WT Parkin compared to overactivation of p-Parkin. Overall, both MALDI-TOF based assays accurately and quantitively measured the E3 ligase activity of Parkin and p-Parkin and the rate at which the co-factor p-Ub-His activates WT Parkin and may further activate p-Parkin.

**Figure 2.**
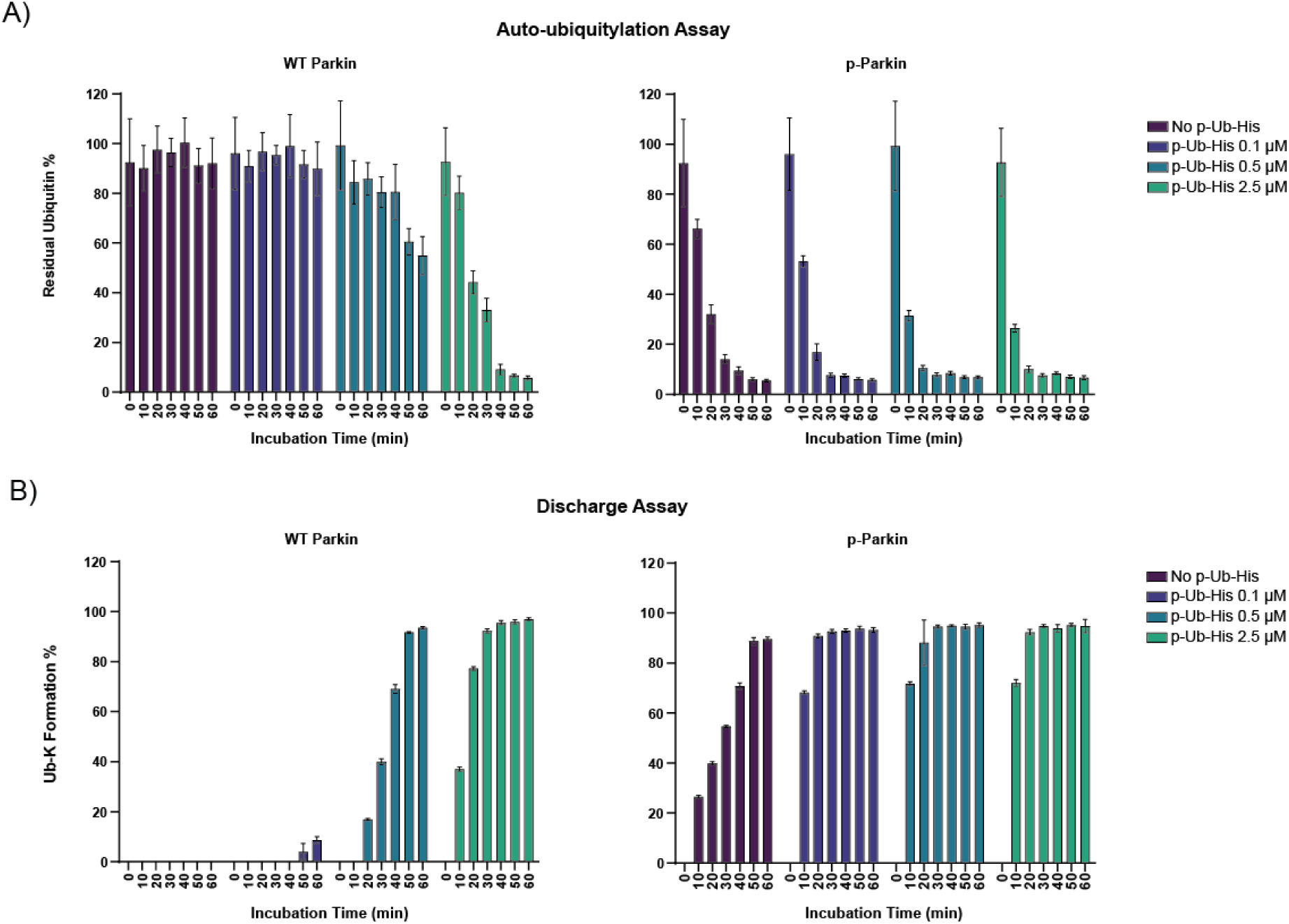
Quantification of WT Parkin and p-Parkin activity by autoubiquitylation (A) and discharge assay (B). WT Parkin and p-Parkin were incubated in absence or in presence of increasing amount of p-Ub-His for up to 60 minutes. The reduction of mono-ubiquitin as consequence WT Parkin and p-Parkin activity is reported as Residual Activity % (A) in autoubiquitylation assay while the Ub-K % formation indicates activity in the discharge assay read-out (B). Data points are reported as the average of 3 replicates ± SD.

**Figure 3.**
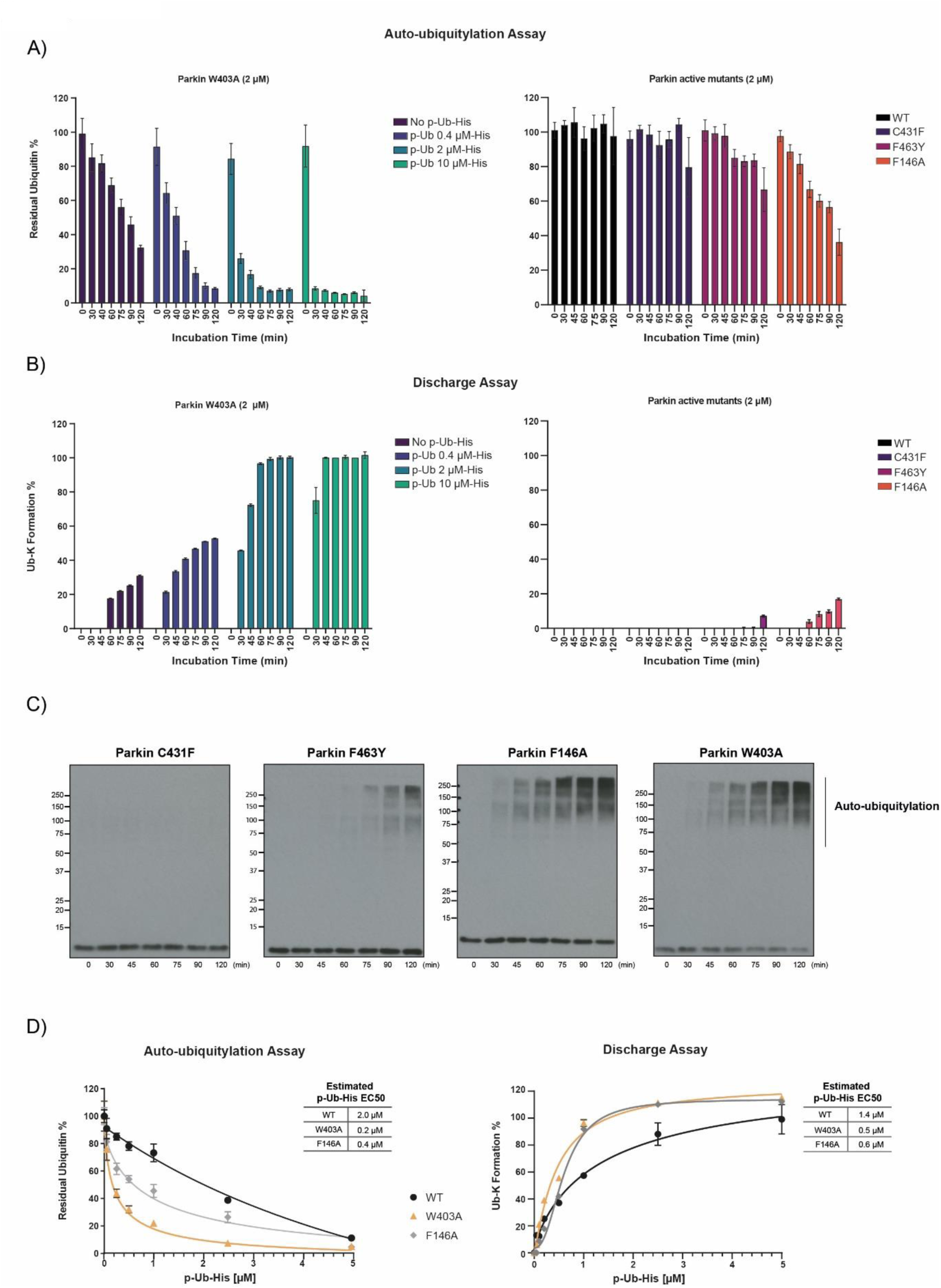
Quantification of Parkin mutants’ activity by autoubiquitylation (A) and discharge assay (B). All Parkin mutants were tested at 2000 nM final at the indicated time points. Results were validated by Parkin *in vitro* ubiquitylation assay followed by SDS-page and western blotting using anti-ubiquitin antibody (C). Estimated Half maximal effective concentration of p-Ub-His for the activation of WT, W430A and F146A Parkin (D). Data points are reported as the average of 3 replicates ± SD

### Quantifying the effect of point mutations on Parkin activity

Structural analysis of inactive and active Parkin has identified three major inter-domain interfaces that that maintain auto-inhibition of Parkin Ub ligase activity^9,10,19^. Based on these studies, engineering point mutations that disrupt the repressor element of Parkin (REP) domain interaction with the RING1 (really interesting new gene 1) domain-interface, W403A, or the RING0-RING2 interface, F146A and F463Y, that each loosen the auto-inhibitory conformation of Parkin and are effective at releasing Parkin activity^9,10,20,21^ as well as rescuing defects in p-Ub binding and Ser65 phosphorylation^21^. We therefore expressed Parkin W403A, F463Y and F146A mutants as well as the catalytic inactive C431A mutant (Sup. Fig. 3C) and compared the impact of these mutations on Parkin activation using both the Parkin autoubiquitylation and discharge MALDI-TOF MS based assays. Since these Parkin activating-mutants only partially release E3 ligase activity, enzymatic concentrations and incubation times were optimized and we consequently observed activity levels lower than activated WT Parkin or p-Parkin. The activity of W403A Parkin could not be detected in absence of p-Ub-His at the concentration of 500 nM (Sup Fig.3) while it was detected at the final concentration of 2 µM (Fig. 3A). The lack of W403A activity at low concentration (500 nM) and in absence of the co-factor p-Ub-His is due to the relatively low level of activity released by this point mutation. Therefore W403A, F146A and F463Y mutants were tested at the final concentration of 2 µM and incubation time extended up to 120 minutes. W403A background autoubiquitylation activity (in absence of p-Ub-His) halves the initial ubiquitin pool in about 90 minutes of incubation (Fig. 3A) and down to a 32.3% of the total at the final time point of 120 minutes. Further activation is achieved in presence of increasing amounts of p-Ub-His (Fig. 3A). The F146A mutant showed a level of activity comparable to those of W403A, with only 36.2% of the initial pool of ubiquitin still detectable after 120 minutes. The active mutant F463Y was about 50% less active compared to W403A and F146A mutants (66.7% of ubiquitin still present after 120 minutes). A similar trend was observed in the discharge assay, albeit the measured activity of the mutants in the discharge assay were relativity low that likely reflects the distinct temperature at which respective assays were performed. We confirmed these findings using an orthogonal Parkin *in vitro* assay in which mutant Parkin F463Y, F164A and W403A and C431F were incubated in the presence of adenosine triphosphate (ATP), MgCl_2,_ E1 ubiquitin-activating ligase, UbcH7 conjugating E2 ligase and ubiquitin. After 60 minutes, reactions were terminated with SDS sample buffer in the presence of 2-mercaptoethanol and heated at 100 ºC, and ubiquitylation was assessed by immunoblot analysis with antibodies that detect ubiquitin (Fig 3C). We further employed the MALDI-TOF based assays to estimate the half maximal effective concentration (EC50) of p-Ub-His for the activation of WT, W403A and F126A Parkin. We incubated 500 nM WT, W430A and F146A Parkin with increasing concentrations of p-Ub-His (0.05 µM, 0.2 µM, 0.5 µM, 1 µM, 2.5 µM and 5 µM) and incubated the reaction 30 minutes in the previously defined conditions. An estimated EC50 of 2 µM for WT Parkin, 0.2 µM for W403A and 0.4 µM for F146A was determined in the MALDI-TOF autoubiquitylation assay settings. Similar trend was observed for the MALDI-TOF discharge assay: 1.4 µM for WT Parkin, 0.5 µM for W403A and 0.6 µM for F146A. The results confirmed that both W403A and F146A Parkin mutants require reduced amount of p-Ub-His to achieve activity levels comparable to those of WT Parkin. Both assays indicates that the W403A mutation requires between 2 and 10 less p-Ub-His to achieve WT Parkin activity levels. Overall, our analysis of Parkin mutants is consistent with the previous literature reporting W403A as one of the most activating Parkin single point mutations^9,10,19^. Moreover, the accurate quantification of the absolute and relative activation effect of Parkin point mutations further validates the ability of both MALDI-TOF based assays to identify Parkin activation and inhibition rates.

### Development of Parkin High-Throughput Screen (HTS)

Primary, activity based high-throughput screening (HTS) represents often the first step when starting a new drug discovery project that targets an enzyme. Such a step is fundamental for the identification of promising candidates from the vast number of natural and synthetic compound libraries available. Since PD is caused by loss of function of Parkin, the pharmaceutical intent is to re-instate the enzymatic activity of Parkin through the identification of Parkin-specific activators. The Parkin auto-ubiquitylation assay relies on the progressive reduction of the ubiquitin signal. In such conditions, the identification of activators will be limited by the assay window itself. On the other hand, the discharge assay offers a larger assay window and the possibility to work at lower temperature (25°C). Therefore, we tested the feasibility of employing the MALDI-TOF based discharge assay to perform a preliminary high-throughput screen for the identification of p-Parkin activators. We tested a compound library of about 20000 compounds predicted to be able to permeate the blood brain barrier. The HTS workflow was designed to be scalable and adaptable for a high-throughput screening campaign and consists of 3 steps: pre-incubation of 5 µL enzymatic mixture with compounds (10 µM in 100% DMSO), reaction initiation by adding 5 µL of substrate (mono-ubiquitin and 50 mM Ac-K) and reaction termination with 5 µL 6% TFA (Fig 4A). A total of 60 × 384 well plates were divided into nine smaller batches of up to 8 × 384 well plates (about 2800 compounds) to be processed daily (Fig. 4 D and E). The use of high-density 1536 AnchorChip MALDI targets allowed to combine up to four 384 assay plates into one MALDI-TOF MS run (Fig 4A). Each plate included a column (16 wells) reserved for positive controls (no compound present, only DMSO) and one column for negative controls (reaction in absence of p-Parkin where only background reading should be detected, example data in Fig. 4B). Data were normalized by dividing the area of the substrate (Ub) to the area of the product (Ub-Ac-K). A linearity curve with known amounts of Ub and Ub-Ac-K was interpolated and used to translate Ub-Ac-K /Ub ratio into % of Ub-K formation (Sup. Fig 5). The robustness of HTS screening is a function of both the variability of positive and negative controls and the statistical space for the robust identification of the compound related effect. A Z’ Prime value > 0.5 is considered a robust assay. The Z’ Prime average for the MALDI-TOF discharge assay was 0.75 with only one 384 plate scoring below the threshold of 0.5 (Fig. 4C) confirming the robustness of the assay and the employability in HTS campaigns. An arbitrary and stringent hit cut-off of +/- 25% activity compared to the control was applied to select compounds to be further investigated. A total of 5 compounds reduced p-Parkin activity by more than 25% and only 1 compound scored as potential activator for a total of 6 positive hits. Given the low number of compounds tested, it was not unexpected that none of the positive hits were confirmed by subsequent validation analysis by IC50 calculation. Identification of genuine active compounds, inhibitors and particularly activators, are a few and far in between, however the HTS screening results indicated an exceptionally low false positive rate (FPR) of 0.028% confirming the advantages of MALDI-TOF based read-out compared to fluorescence-based approaches. Several activators of Parkin have been described in patent literature, although peer-reviewed research is not available. We tested three molecules reported in patent WO 2018/023029 (chemotype B1, B2 and C1, Sup. Fig. 6A) for their ability to activate Parkin. All compounds were tested at a final concentration of 50 µM in a time course experiment (7 time points) against WT-Parkin (activated by equimolar amounts of p-Ub-His) and p-Parkin using both the MALDI-TOF autoubiquitylation assay (Sup Fig. 6B) and the discharge assay (Sup. Fig. 6C). The MALDI-TOF autoubiquitylation assay did not detect a statistically significant differences between the control (DMSO only) and the tested compounds against either WT Parkin or p-Parkin (Sup. Fig. 6B). However, the MALDI-TOF discharge assay successfully detected a small but statistically significant activation effect on WT-Parkin in presence of chemotype B2 after 50 and 60 minutes of incubation while no effect was observed against p-Parkin. These results indicate that the MALDI-TOF discharge assay might prove more sensitive for the detection of small changes in enzymatic activity compared to the auto-ubiquitylation assay. This can be explained by the relatively easy access of Ac-Lysine to the Parkin active site already in presence of weak structural perturbations. Overall, our results confirmed the ability of the previously reported chemotype B2 to enhance WT-Parkin and validated the ability of the MALDI-TOF discharge assay to effectively detect Parkin activators.

**Figure 4.**
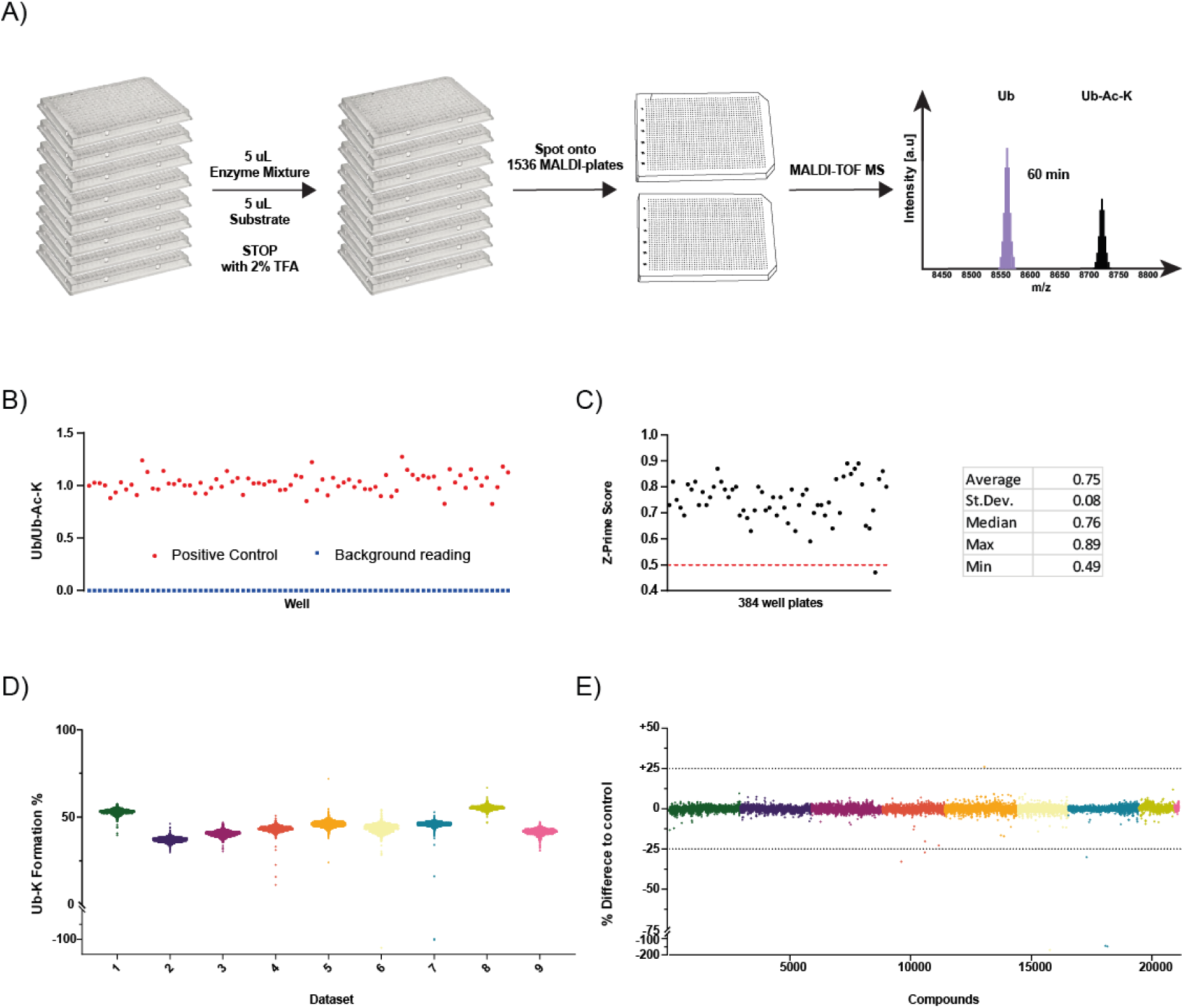
p-Parkin High-Throughput Screening by MALDI-TOF MS discharge assay. Workflow schematic (A) and representative data of positive control and background reading (B). Z’ Prime value for HTS plates (C). Data distribution for independent datasets (experiments were performed in single replicate) and compounded data of HTS normalized to the positive controls for the identification of inhibitors and activators. Compounds have been tested as single replicates while 16 data points were included in each 384 well plate for both positive and negative controls.

## Discussion

Accumulating of biological and structural studies have provided unprecedented understanding of the regulation of Parkin by either direct phosphorylation on serine 65 or by the interaction with phosphorylated ubiquitin. Currently, the *in vitro* quantification of Parkin’s activity relies on the use of SDS-PAGE followed by antibody-based detection of ubiquitylation events. This method enables assessment of Parkin’s activity via monitoring Parkin auto-ubiquitylation pattern; multi-monoubiquitylation of substrates such as MIRO1^22^, mono-ubiquitylation of UBE2L3^10,12^ or the formation of free ubiquitin chains ^10,23,1^. Such approaches are intrinsically low throughput and time consuming. Here we reported two robust MALDI-TOF based assays to investigate the activity of Parkin in a fast, quantitative and high-throughput fashion. Both MALDI-TOF based assays accurately and quantitatively recapitulate Parkin activity and activation rates in presence of the activating cofactor p-Ub-His. Structural studies have revealed point mutations known to partially release Parkin auto-inhibitory state and release background activity. The MALDI-TOF based technology enabled facile comparison and quantification of the relative impact of such point mutations on Parkin activity. Whilst this assay will aid in understanding the regulation of Parkin activity by academic researchers, MALDI-TOF based technologies are emerging as the gold standard in the drug discovery space. Fluorescence probes as UbMes and UbFluor have been reported as functional Ub-based probes for determining Parkin activity^*15*^. Such strategies are potentially scalable to high-throughput screening levels; however, the use of fluorescence as analytical read-out is inherently problematic because of fluorescence artefacts that result in both false positives and false negatives. For example, UbFluor is labile in the presence of reducing agents or other small molecules that possess thiol or amine groups that may cleave UbFluor even in the absence of RBR E3, resulting in false positives^14^. Fluorescent small molecules may also disrupt fluorescence polarization readings, resulting in false negatives. Rate of false positive and negative is highly dependent on the fluorophore used, the stability of the substrate, the assay conditions and the nature of the chemical libraries tested. A recent study suggested that false discovery rate might score anywhere between 0.5 to 9.9% depending on the assay and type of fluorescence used^24^. This translates into the risk of following up on false leads, with obvious consequences in terms of increased costs and reduced efficiency. The screening of ∼20000 compounds by MALDI-TOF MS discharge assay resulted in a false positive rate of only 0.028%, well below what to be expected with fluorescent based tools.

The social and economic impact of PD has sustained intense research efforts to identify pharmacological treatments, producing several patents reporting chemical structures of Parkin activators. Here we tested three previously reported molecules using both the MALDI-TOF MS auto-ubiquitylation and discharge assay. Our results confirmed the expected Parkin-activation effect of one of these molecules by the MALDI-TOF discharge assay while no activation effect was observed by the MALDI-TOF auto-ubiquitylation assay. Notably, the HTS MALDI-TOF base strategy can also be easily applied to other RBR E3 ligases (as has been done for HOIP^16^) including those E3 ligases that peculiarly discharge on non-canonical residues (for example serine and sugars) such as HOIL-1 and RNF213^25,26^. Overall, we anticipate MALDI-TOF based technologies to substantially increase our understanding of the functioning of E2 conjugating enzymes and E3 ligases by providing accurate and quantitative data and to contribute to drug discovery campaigns in the ubiquitin field.

## Materials & Methods

### Reagents

Ubiquitin was acquired by sigma Aldrich (U6253). Eppendorf Low-Bind 384 well plates (Cat. Number 951031305) were used for low-throughput assay. Ac-K was purchased from Bachem/Cambridge (Cat. Number 4000486.0001). p-Ub-His, ^15^N Ubiquitin, Ube1, Ube2L3, WT and phosphorylated Parkin and Parkin point mutants were expressed and purified in house as indicated in below.

### Autoubiquitylation MALDI-TOF MS Parkin Activity assay

200 nM E1 activating enzyme, 1000 nM UBE2L3 conjugating enzyme, 1000 nM WT parkin or p-Parkin, 20 mM MgCl_2_, 2 mM ATP, 0.05% BSA and 2 mM TCEP were mixed in 1X phosphate buffer (PBS, pH 8.5) and aliquoted into Eppendorf Low-Bind plates (5 µL per well). The reactions were started by adding 5 µL of 50 µM Ubiquitin (in 1X PBS, pH 8.5) supplemented with the indicated amount of p-Ub-His. Plates were sealed with adhesive aluminium foil and incubated at 37°C in an Eppendorf ThermoMixer C (Eppendorf) equipped with a ThermoTop and a SmartBlock™ PCR 384. The reactions were stopped at the indicated time points by the addition of 5 µL 6% TFA supplemented with 6 µM ^15^N Ubiquitin. Samples were spotted on 1536 AnchorChip MALDI target using a Mosquito nanoliter pipetting system (TTP Labtech) and analysed by MALDI-TOF MS as previously reported^16^.

### Discharge MALDI-TOF MS Parkin Activity assay

An identical enzymatic mixture as the autoubiquitylation assay was prepared. The reactions were started by adding 5 µL of 50 µM Ubiquitin supplemented with the indicated amount of p-Ub-His and 50 mM Ac-K. Plates were incubated at room temperature (25 degrees) and sealed with adhesive aluminium foil. The reactions were stopped at the indicated time points by the addition of 5 µL 6% TFA.

### Parkin HTS screening

All Parkin HTS assays were performed in a total volume of 20.01µl at room temp using a FluidX Xrd-384 dispenser. To plates containing 20nl of compound 10ul of a mix containing 500nM p-PARKIN, 400nM UBE1, 4000nM UBE2L3, 20µM MgCl2, 2mM ATP in a 50mM HEPES pH8.5 20mM TECEP buffer was added. The plates were preincubated at 25°C for 30mins and the assay was then initiated with the addition of 10µL of Ubiquitin mix containing 100µM Ubiquitin, 100mM Ac-lysine. The assay was incubated for 20mins at 25°C. The assay was then terminated with the addition of 10µl 6% TFA.

### Expression and Purification of recombinant GST-PINK1 126-end (pediculus humanus)

BL21 codon plus cells were transformed with MRC-PPU plasmid DU34798. A single antibiotic resistant colony was selected and propagated for 16 h at 37°C, 200 rpm. 12 × 1L batches of LB broth/carbenicillin were inoculated with the overnight culture and grown until an OD_600_ of 0.8. The incubation temperature was dropped to 26°C and PINK1 expression was induced by supplementing the media with 0.1 mM Isopropyl β-D-1-thiogalactopyranoside (IPTG) and left to express for overnight. The cells were collected by centrifugation (25 min at 4200 rpm) and the clarified broth was decanted. The cells were resuspended in 20 ml per pellet of 50 mM Tris pH 7.5, 250 mM NaCl, 1 mM DTT, 1 mM AEBSF, 10 µg/ml Leupeptin. The suspension was collected into 50 ml centrifuge vials, chilled on ice and sonicated using 6 pulses of 55% amplitude and 15 s pulses. The suspension was clarified by centrifugation at 40000 x g for 25 min at 4°C. 6 ml GSH-agarose was equilibrated with wash buffer (50 mM Tris pH 7.5, 250 mM NaCl, 1 mM DTT) and mixed with the clarified cell lysate for 90 min. The GSH-agarose was recovered by sedimentation, washed 5 times with 5 volumes of wash buffer and eluted in wash buffer containing 10 mM reduced GSH.

### Expression and Purification of recombinant Parkin 1-465 (human), Parkin active mutants and p-Parkin

Human wild type Parkin 1-465 along with the F146A, W403A, and F463Y mutants (MRC-PPU plasmids DU40847, DU44642, DU44643 and DU58844) were expressed as His6-SUMO-fusion proteins and purified as described previously^12^

To produce phosphorylated Parkin, the fusion protein was captured on Ni-agarose, washed and incubated with 5 mg of GST-PINK1 126-end in the presence of 10 mM MgCl_2_ and 2 mM ATP for 4 h at 27°C. The initial kinase and Mg-ATP were removed and replaced with fresh kinase and Mg-ATP for incubation over night at 27°C. The Ni-agarose was washed three times with wash buffer and Parkin was eluted in the smallest possible volume. The protein was then dialysed in the presence of SENP1 as previously described^12,13^ The protein was further phosphorylated with more PINK1 and Mg-ATP and at the same time concentrated to 6 mg/ml. Finally, the protein was purified further by chromatography on a Superdex 200 as described above and concentrated to about 2 mg /ml. Note that phosphorylated Parkin is more soluble than unphosphorylated Parkin and yields are generally higher.

### Expression and Purification of recombinant p-Ubiquitin-His (pSer65-Ubiquitin-6His), ^15^N Ubiquitin and Ub-K

Ubiquitin-His_6_ was produced from a kanamycin resistance conferring plasmid MRC-PPU reagent DU21990. The cells were grown and induced as for untagged ubiquitin, but they were collected and lysed in 50 mM Tris pH 7.5, 250 mM NaCl, 25 mM imidazole, 7 mM 2-mercaptoethanol, 10 µg/ml Leupeptin (Apollo Scientific), 1 mM AEBSF (Apollo Scientific). The protein was purified over Ni-NTA agarose, eluted into a 0.4 M imidazole buffer and dialysed against 50 mM Tris pH 7.5, 200 mM Tris pH 7.5, 7 mM 2-mercapto ethanol. For phosphorylation at Ser65, 20 mg of Ubiquitin-His was incubated with 2 mg of GST-PINK1 in the presence of 10 mM MgCl_2_ and 2 mM ATP for overnight at 28°C. The Ubiquitin-His was collected on 1 ml Ni-NTA agarose, washed 4 times with 12 bed volumes of 50 mM Tris pH 7.5, 200 mM Tris pH 7.5, 7 mM 2-mercapto ethanol and recovered by elution with imidazole. Imidazole was removed and p-Ub-His concentrated using Millipore Ultra filter (3000 MWCO) followed by subsequent sample dilution in 1x PBS, pH 7.0. The sequence was repeat for 6 times using a 6-fold dilution. Phosphorylation efficiency was assessed by LC-MS analysis (Sup. Fig 1): 70% of Ub-His was successfully phosphorylated. No further purification step was performed, the relative purity was considered in the experimental calculations. Expression and purification of ^15^N-Ubiquitin and Ub-K was performed as previously reported^27,28^.

M.M.K.M. is supported by a Wellcome Trust Senior Research Fellowship in Clinical Science (210753/Z/18/Z); the Michael J Fox Foundation; and an EMBO YIP Award.

---

**Sup Figure 1.**
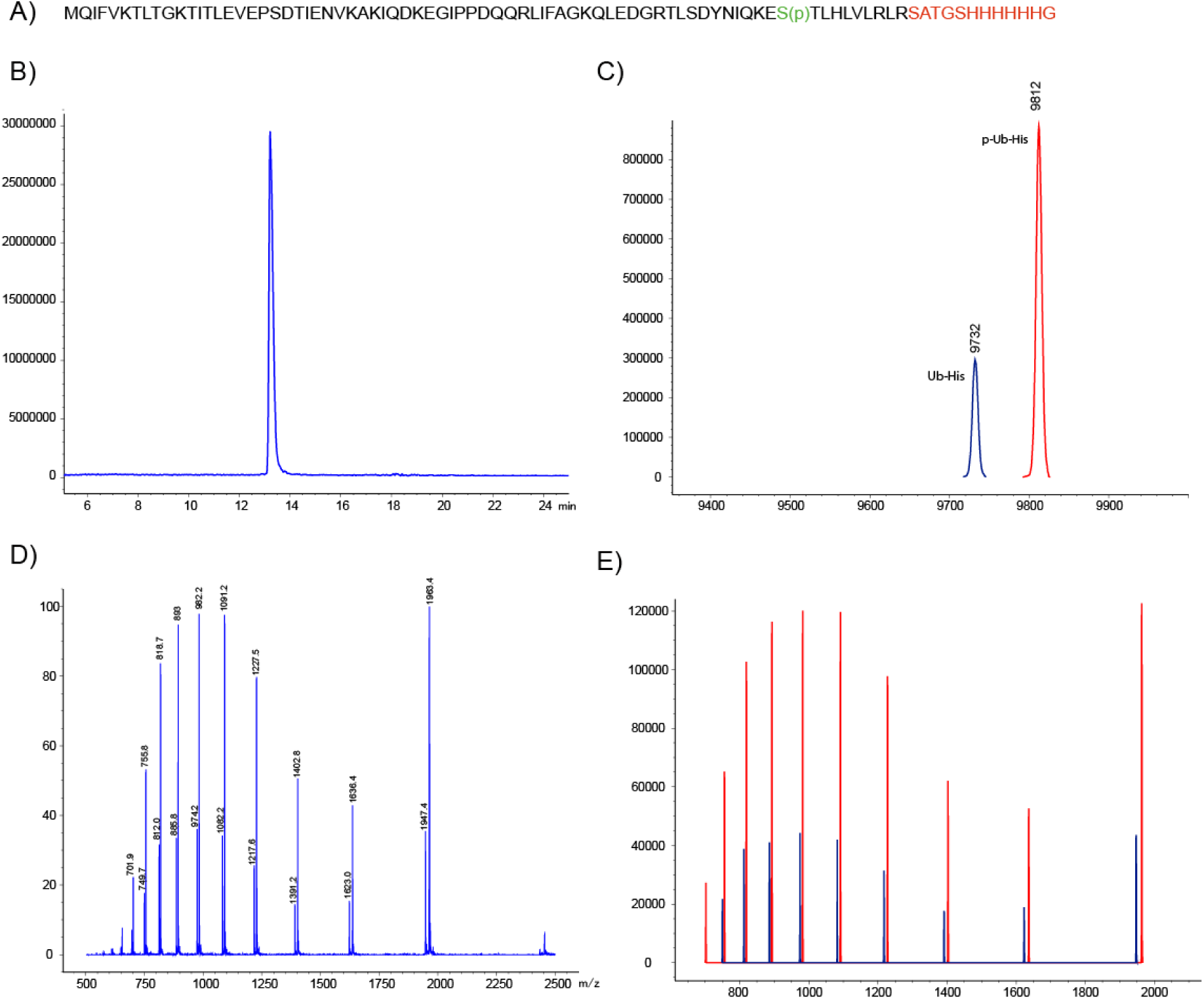
His tagged Phosphorylated Ubiquitin (p-Ub-His) Quality Control. p-Ub-His was expressed and purified as indicated in Methods. A) Protein sequence highlighted in green Serine 65 and in red 6His tag sequence. B) Chromatogram and C) MS components: 9812 m/z corresponding to the p-Ub-His expected m/z and 9732 m/z corresponding to the remaining not phosphorylated counterpart. Purity level have been considered into experimental calculations. D) Mass Spectrum and E) Deconvoluted Ion Set. p-Ub-His estimated at 70%.

**Sup. Figure 2.**
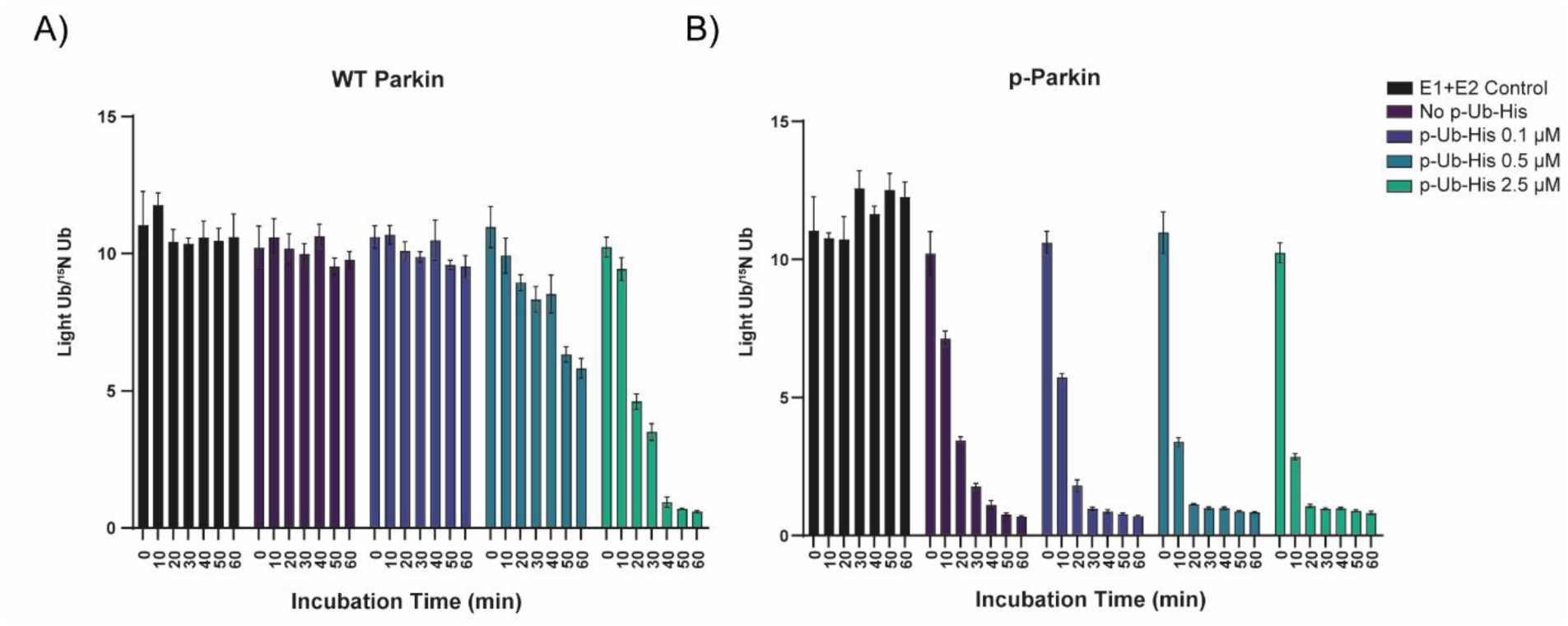
Light/^15^N ubiquitin ratio before conversion into Remaining Ubiquitin %. Stable level of light/^15^N ubiquitin in the E1+E2 control indicate no consumption of ubiquitin in presence of the E1 activating enzyme and UBE2L3 conjugating enzyme only.

**Sup. Figure 3.**
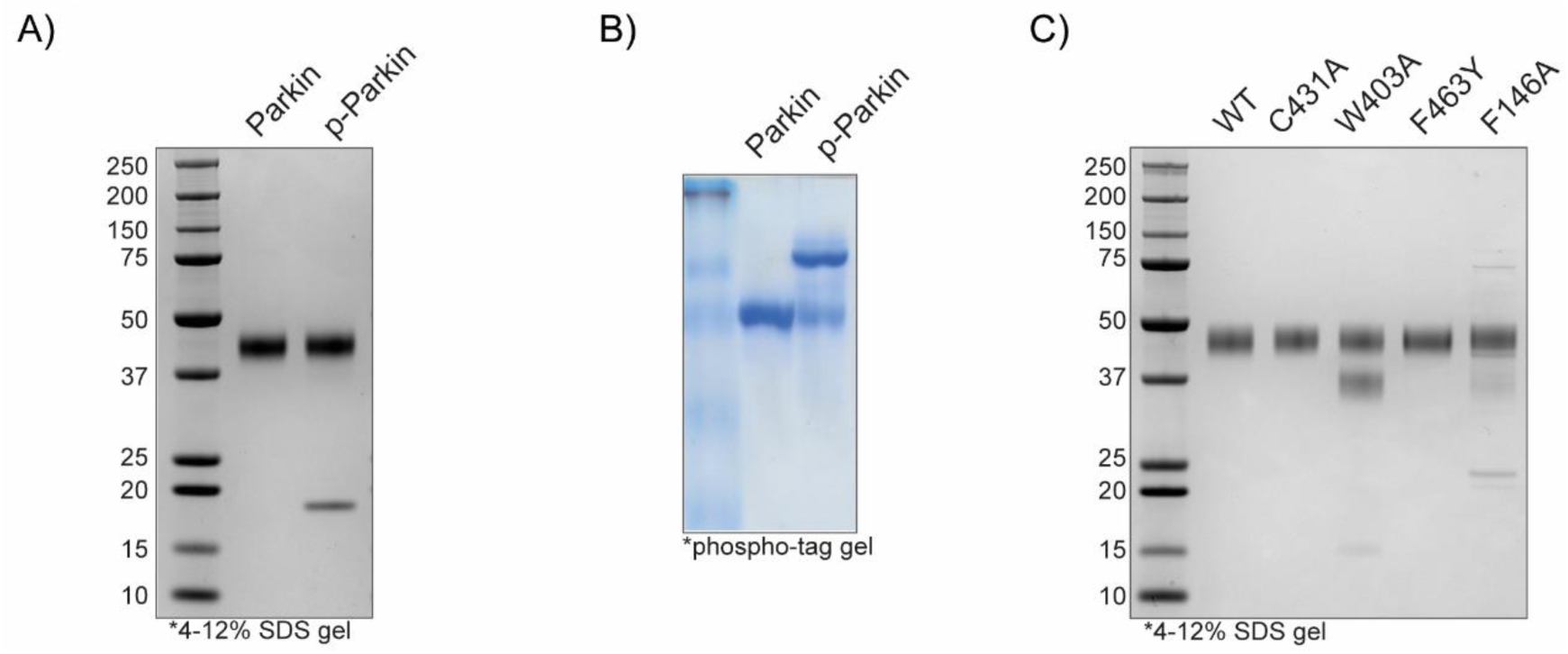
Parkin, p-Parkin and activating Parkin mutants. **A)** Parkin and p-Parkin purity check by SDS-page. B) Parkin phosphorylation efficiency, phosphor-tag gels indicates that p-Parkin is about 70% pure. C) Parkin activating mutants purity check by SDS-page.

**Sup. Figure 4.**
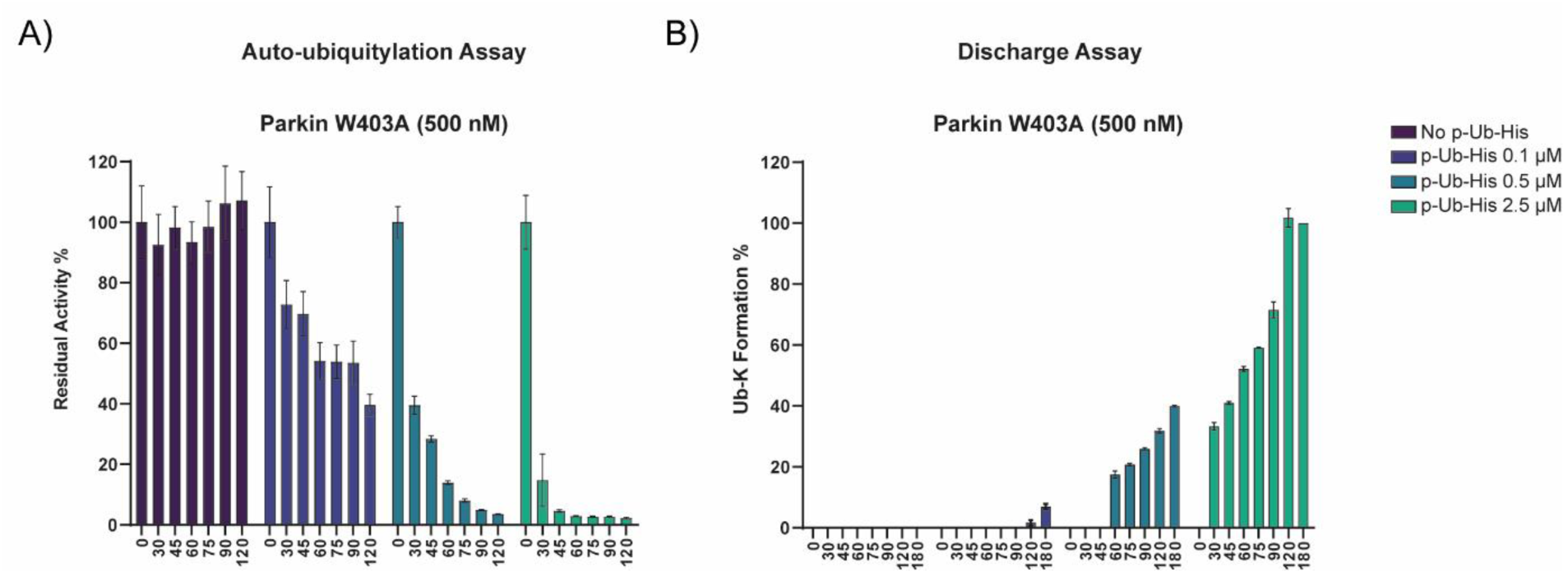
Parkin W403A autoubiquitylation (A) and discharge assay (B) at 500 nM. No background activity was detected when testing W403A at the final concentration of 500 nM.

**Sup. Figure 5.**
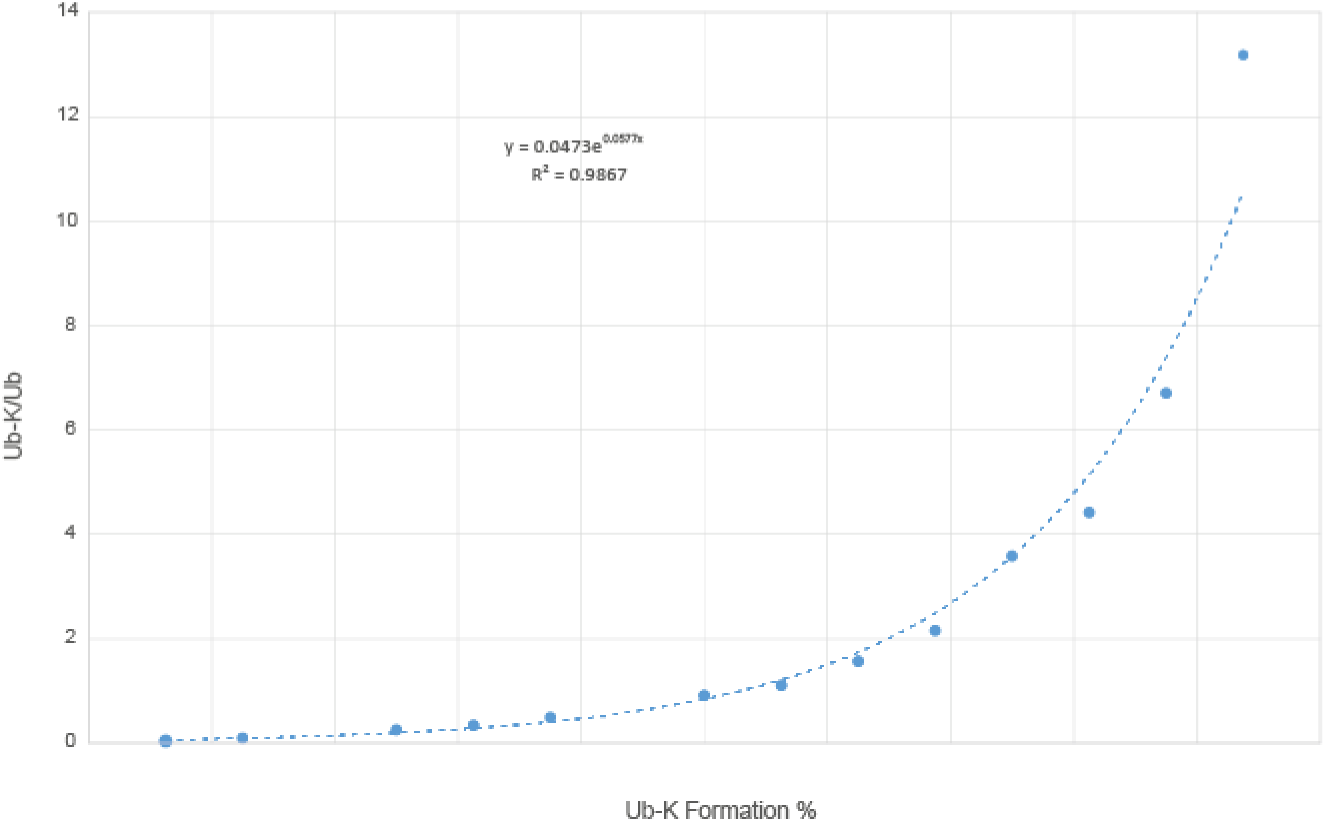
Linearity Curve for Ub-K formation %. Known amount of Ub (substrate) and Ub-K (product) were mixed and analysed by MALDI-TOF MS. Resulting curve and associated exponential equation was employed to translate Ub-K/Ub peak area ratio into Ub-K Formation%

**Sup. Fig. 6.**
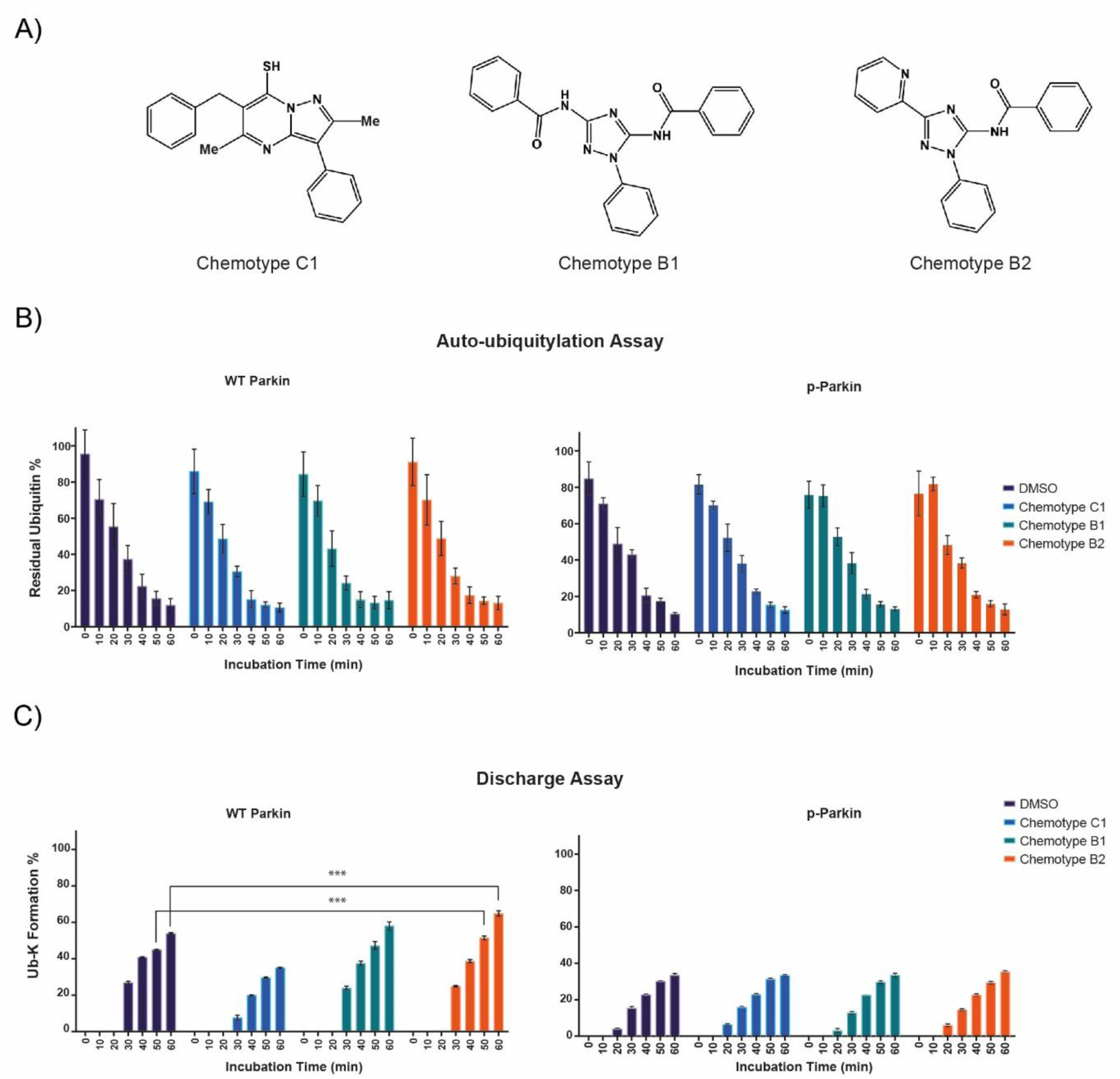
Evaluation of previously reported Parkin activators. Three compounds selected from patent WO 2018/023029 (panel A: chemotype B1, B2 and C1) were tested against WT-Parkin and p-Parkin at the final concentration of 50 µM by autoubiquitylation (B) and discharge assay (C). Reactions were stopped at the indicated time points. Data points are reported as the average of 2 replicates spotted in duplicate (4 data points total) ± SD.

## References

1 Corti, O., Lesage, S. & Brice, A. What genetics tells us about the causes and mechanisms of Parkinson’s disease. Physiol Rev 91, 1161–1218, doi:10.1152/physrev.00022.2010 (2011).

2 Kitada, T. et al. Mutations in the parkin gene cause autosomal recessive juvenile parkinsonism. Nature 392, 605–608, doi:10.1038/33416 (1998).

3 Valente, E. M. et al. Hereditary early-onset Parkinson’s disease caused by mutations in PINK1. Science 304, 1158–1160, doi:10.1126/science.1096284 (2004).

4 Miller, S. & Muqit, M. M. K. Therapeutic approaches to enhance PINK1/Parkin mediated mitophagy for the treatment of Parkinson’s disease. Neurosci Lett 705, 7–13, doi:10.1016/j.neulet.2019.04.029 (2019).

5 Narendra, D. P. & Youle, R. J. Targeting mitochondrial dysfunction: role for PINK1 and Parkin in mitochondrial quality control. Antioxid Redox Signal 14, 1929–1938, doi:10.1089/ars.2010.3799 (2011).

6 Koyano, F. et al. Ubiquitin is phosphorylated by PINK1 to activate parkin. Nature 510, 162–166, doi:10.1038/nature13392 (2014).

7 Shiba-Fukushima, K. et al. PINK1-mediated phosphorylation of the Parkin ubiquitin-like domain primes mitochondrial translocation of Parkin and regulates mitophagy. Sci Rep 2, 1002, doi:10.1038/srep01002 (2012).

8 Chaugule, V. K. et al. Autoregulation of Parkin activity through its ubiquitin-like domain. EMBO J 30, 2853–2867, doi:10.1038/emboj.2011.204 (2011).

9 Kumar, A. et al. Disruption of the autoinhibited state primes the E3 ligase parkin for activation and catalysis. EMBO J 34, 2506–2521, doi:10.15252/embj.201592337 (2015).

10 Trempe, J. F. et al. Structure of parkin reveals mechanisms for ubiquitin ligase activation. Science 340, 1451–1455, doi:10.1126/science.1237908 (2013).

11 Sauve, V. et al. A Ubl/ubiquitin switch in the activation of Parkin. EMBO J 34, 2492–2505, doi:10.15252/embj.201592237 (2015).

12 Kondapalli, C. et al. PINK1 is activated by mitochondrial membrane potential depolarization and stimulates Parkin E3 ligase activity by phosphorylating Serine 65. Open Biol 2, 120080, doi:10.1098/rsob.120080 (2012).

13 Kazlauskaite, A. et al. Parkin is activated by PINK1-dependent phosphorylation of ubiquitin at Ser65. Biochem J 460, 127–139, doi:10.1042/BJ20140334 (2014).

14 Foote, P. K. & Statsyuk, A. V. Monitoring PARKIN RBR Ubiquitin Ligase Activation States with UbFluor. Curr Protoc Chem Biol 10, e45, doi:10.1002/cpch.45 (2018).

15 Park, S., Foote, P. K., Krist, D. T., Rice, S. E. & Statsyuk, A. V. UbMES and UbFluor: Novel probes for ring-between-ring (RBR) E3 ubiquitin ligase PARKIN. J Biol Chem 292, 16539–16553, doi:10.1074/jbc.M116.773200 (2017).

16 De Cesare, V. et al. The MALDI-TOF E2/E3 Ligase Assay as Universal Tool for Drug Discovery in the Ubiquitin Pathway. Cell Chem Biol 25, 1117–1127 e1114, doi:10.1016/j.chembiol.2018.06.004 (2018).

17 Wenzel, D. M., Lissounov, A., Brzovic, P. S. & Klevit, R. E. UBCH7 reactivity profile reveals parkin and HHARI to be RING/HECT hybrids. Nature 474, 105–108, doi:10.1038/nature09966 (2011).

18 Kazlauskaite, A. et al. Binding to serine 65-phosphorylated ubiquitin primes Parkin for optimal PINK1-dependent phosphorylation and activation. EMBO Rep 16, 939–954, doi:10.15252/embr.201540352 (2015).

19 Wauer, T. & Komander, D. Structure of the human Parkin ligase domain in an autoinhibited state. EMBO J 32, 2099–2112, doi:10.1038/emboj.2013.125 (2013).

20 Riley, B. E. et al. Structure and function of Parkin E3 ubiquitin ligase reveals aspects of RING and HECT ligases. Nat Commun 4, 1982, doi:10.1038/ncomms2982 (2013).

21 Tang, M. Y. et al. Structure-guided mutagenesis reveals a hierarchical mechanism of Parkin activation. Nat Commun 8, 14697, doi:10.1038/ncomms14697 (2017).

22 Kazlauskaite, A. et al. Phosphorylation of Parkin at Serine65 is essential for activation: elaboration of a Miro1 substrate-based assay of Parkin E3 ligase activity. Open Biol 4, 130213, doi:10.1098/rsob.130213 (2014).

23 Antico, O. et al. Global ubiquitylation analysis of mitochondria in primary neurons identifies endogenous Parkin targets following activation of PINK1. Sci Adv 7, eabj0722, doi:10.1126/sciadv.abj0722 (2021).

24 Borrel, A. et al. High-Throughput Screening to Predict Chemical-Assay Interference. Sci Rep 10, 3986, doi:10.1038/s41598-020-60747-3 (2020).

25 Kelsall, I. R., Zhang, J., Knebel, A., Arthur, J. S. C. & Cohen, P. The E3 ligase HOIL-1 catalyses ester bond formation between ubiquitin and components of the Myddosome in mammalian cells. Proc Natl Acad Sci U S A 116, 13293–13298, doi:10.1073/pnas.1905873116 (2019).

26 Otten, E. G. et al. Ubiquitylation of lipopolysaccharide by RNF213 during bacterial infection. Nature 594, 111–116, doi:10.1038/s41586-021-03566-4 (2021).

27 Ritorto, M. S. et al. Screening of DUB activity and specificity by MALDI-TOF mass spectrometry. Nat Commun 5, 4763, doi:10.1038/ncomms5763 (2014).

28 De Cesare, V. et al. Deubiquitinating enzyme amino acid profiling reveals a class of ubiquitin esterases. Proc Natl Acad Sci U S A 118, doi:10.1073/pnas.2006947118 (2021).

